# Development of a new DHFR-based destabilizing domain with enhanced basal turnover and applicability in mammalian systems

**DOI:** 10.1101/2022.06.21.495152

**Authors:** Emi Nakahara, Vishruth Mullapudi, Lukasz A. Joachimiak, John D. Hulleman

## Abstract

Destabilizing domains (DDs) are an attractive strategy allowing for positive post-transcriptional small molecule-regulatable control of a fusion protein’s abundance. Yet in many instances, the currently available DDs suffer from higher-than-desirable basal levels of the fusion protein. Accordingly, we redesigned the E. coli dihydrofolate reductase (ecDHFR) DD by introducing a library of ~1200 random ecDHFR mutants fused to YFP into CHO cells. Following successive rounds of FACS sorting, we identified six new ecDHFR DD clones with significantly enhanced proteasomal turnover in the absence of a stabilizing ligand, trimethoprim (TMP). One of these clones, designated as ‘C12’, contained four unique missense mutations (W74R/T113S/E120D/Q146L) and demonstrated a significant 2.9-fold reduction in basal levels compared to the conventional ecDHFR DD YFP. This domain was similarly responsive to TMP with respect to dose-response and maximal stabilization, indicating an overall enhanced dynamic range. Interestingly, both computational and wet-lab experiments identified the W74R and T113S mutations of C12 as the main contributors towards its basal destabilization. Yet, the combination of all the C12 mutations were required to maintain both its enhanced degradation and TMP stabilization. We further demonstrate the utility of C12 by fusing it to IκBα and Nrf2, two stress-responsive proteins that have previously been challenging to regulate. In both instances, C12 significantly enhanced the basal turnover of these proteins and improved the dynamic range of regulation post stabilizer addition. These advantageous features of the C12 ecDHFR DD variant highlight its potential for replacing the conventional N-terminal ecDHFR DD, and overall improving the use of destabilizing domains, not only as a chemical biology tool, but for gene therapy avenues as well.

## INTRODUCTION

The inducible synthesis of proteins is an important fundamental ability of all cells to respond to various stimuli and stresses. Likewise, the regulated control of protein therapeutics not only serves as a common/essential strategy for cell biology research^1^, but also as a potential avenue for more nuanced gene therapy applications. Currently available gene therapy approaches rely on the delivery of a constitutively-expressed transgene, which are primarily appropriate for loss-of-function disorders. Yet, given the number of more complex diseases, as well as instances wherein gene therapies could lead to potential off-target or deleterious effects (phenotoxicity)^2^, it is important to develop or optimize existing methods for conditionally regulating such approaches. Destabilizing domains (DD), which enable inducible protein control via the administration of a small molecule ligand^3^, offer a promising solution to the conditional regulation challenge. DDs are engineered protein domains that are targeted for proteasomal degradation upon translation in cells; however, addition of a small molecule pharmacological chaperone which binds to and stabilizes the DD, prevents its degradation and promotes its abundance within the cell. These DDs can serve as an appendable element to a protein of interest, conferring its destabilizing properties.

Since the original development of the FK506 binding protein (FKBP)-derived DD and Shield 1 stabilizing ligand system^3^, multiple new DDs and stabilizers have also been generated with varying biochemical characteristics that suit different experimental purposes and/or model systems^4–7^. The E. coli dihydrofolate reductase (ecDHFR) based DD, which is stabilized by the antibiotic trimethoprim (TMP), is of particular interest due to its sensitivity to low concentrations of TMP, as well as the widespread availability and *in vivo* stability of TMP^4^. Together, the FKBP and ecDHFR based DD systems have been successfully used in a variety of applications, from stress responsive signaling^8–10^ to neuroprotection^11^.

However, as with many first-generation regulatory systems, there are aspects of the DD system that could be improved upon, including optimization of ‘leaky’ expression under basal conditions. For example, when used in transient transfection experiments, and in some instances of stable cells, the ecDHFR DD also suffers from less-than-ideal basal turnover rates^12–14^. For example, human embryonic kidney (HEK-293A) cells transiently overexpressing ecDHFR DD.YFP exhibit a considerable amount of punctate fluorescence without the addition of TMP^15^. While the ecDHFR DD is not technically stabilized under these conditions (it actually appears to reside in aggresomes^15^), it nonetheless builds up and can lead to unwonted (and uncontrolled) basal expression. Other groups which have utilized the ecDHFR DD system have run into similar issues of leaky expression^13^, therefore requiring either multiple DDs to be attached to a protein of interest^16^, or another orthogonal regulatory system such as Tet-ON/OFF to be applied in conjunction with the DD system^14^.

Herein, we sought to improve ecDHFR DD characteristics by generating and screening for new ecDHFR mutants that have improved basal degradation without sacrificing TMP-induced stabilization. We discovered six new and unique ecDHFR mutants with significantly improved basal degradation, two of which were able to be fully stabilized by TMP. After narrowing our focus to a novel ecDHFR DD dubbed as ‘C12’, we performed computational stability analysis, site-directed mutagenesis, and truncation studies to further characterize the molecular requirements for this domain. Lastly, we fused C12 to two previously recalcitrant proteins, inhibitor of nuclear factor kappa B (IκBα) and nuclear-erythroid factor 2-related factor 2 (Nrf2), demonstrating significantly enhanced basal turnover of these client proteins, increased dynamic range, and control of downstream signaling. The culmination of these results suggests that researchers should strongly consider utilizing the C12 ecDHFR DD as a replacement for the conventionally used N-terminal ecDHFR DD in future studies.

## RESULTS

### Generation and screening of a new ecDHFR mutant library reveals a set of TMP-stabilized ecDHFR DDs with unique mutations

An inherent challenge of developing conditional drug-controlled transcriptional/translational regulation systems is maintaining low basal levels of activity in the absence of the small molecule regulator. To improve upon the currently available ecDHFR DDs, we performed error-prone PCR on the wildtype (WT) ecDHFR sequence to generate a library of ~1,200 mutant genes containing 8-10 mutations/kb (Fig. 1). To easily track the degradation and stabilization of the newly developed ecDHFR mutants, we fused a C-terminal YFP reporter to the end of each ecDHFR mutant and then transduced this library into CHO cells (multiplicity of infection ~ 1, Fig. 1). This parental population was processed by fluorescence-activated cell sorting (FACS) twice to isolate a population of cells lacking detectable fluorescence (i.e., a ‘dark’ population). The dark population was then treated with TMP (10 μM, 24 h) before a final round of sorting, this time isolating single-cell clones with fluorescence into 96 well-plates (Fig. S1).

**Figure 1.**
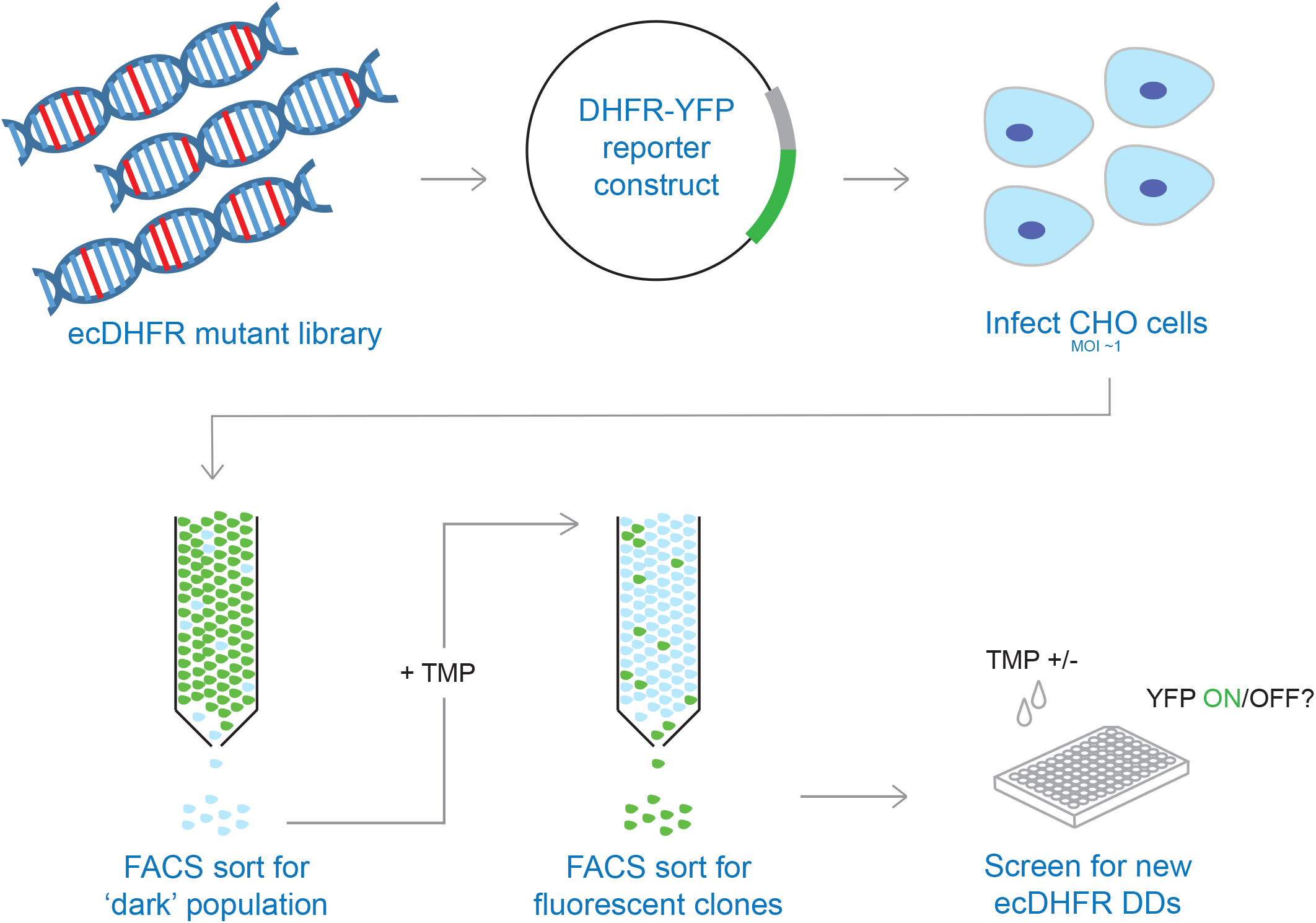
Schematic of screening process for new E. coli DHFR (ecDHFR) DDs. A ecDHFR mutant library was generated with each mutant fused to YFP and expressed in a CHO cell population that was FACS sorted into TMP-responsive single-cell clones.

Each single-cell clone was assessed for fluorescence under basal and TMP-induced conditions to identify clones with the best basal turnover of the ecDHFR DD and dynamic range after stabilization. Following these screening steps, we identified six clones (denoted as A5, C12, F8, F10, G10, G11) that behaved ideally (i.e., lowest basal fluorescence, highest TMP-induced fluorescence). The ecDHFR DNA from each of these clones was amplified and sequenced from the genomic DNA of the single colony clones, revealing a surprisingly diverse breadth of mutations across the ecDHFR sequence (Table 1). None of the newly-identified mutants share a single common mutation with the conventional N-terminal or C-terminal ecDHFR DD^4^, while the only shared mutation site among three of the six new mutants was W74. These observations suggest that a variety of residues across the ecDHFR domain are important for its stability within mammalian cells, and that the ecDHFR domain tolerates a variety of point mutations while still maintaining its ability to bind to TMP, likely indicating that many residues cooperate together to help bind TMP^17^.

**Table 1.** Mutations of new and conventionally-used ecDHFR DDs.

| **clone** | **mutations** |
| --- | --- |
| A5 | M16K/A26A/T46S/D87G/A117A/V119Q |
| C12 | W74R/T113S/E120D/A143A/Q146L |
| F8 | T68A/R71R/V72D/W74C/V88V |
| F10 | D27D/W74C/D116V/N147S |
| G10 | K38R/E80V/Y128F |
| G11 | S49T/I50F/M92K |
| conv N term | R12Y/G67S/Y100I |
| conv C term | R12H/N18T/A19V/G67S |

### Characterizing new TMP-stabilized ecDHFR DDs with enhanced basal degradation and similar stabilization propensity to the conventional ecDHFR DD

With the discovery of these six new ecDHFR DD mutants in stable CHO cells, we next tested their behavior under more proteasomally-challenging conditions; when transiently overexpressed in cell culture. In this scenario, the conventional ecDHFR DD shows detectable YFP fluorescence even in the absence of stabilizer^15^. We transiently transfected each of the new ecDHFR DDs into HEK-293A cells under conditions where the conventional ecDHFR DD produces obvious basal YFP fluorescence (Fig. 2A). The fluorescence of all six new ecDHFR DD mutants was noticeably reduced compared to the conventional ecDHFR DD under basal conditions (Fig. 2A). Moreover, TMP treatment of the same transfected cells (10 μM, 24 h) resulted in similar levels of stabilization compared to the conventional N-terminal ecDHFR (Fig. 2B).

**Figure 2.**
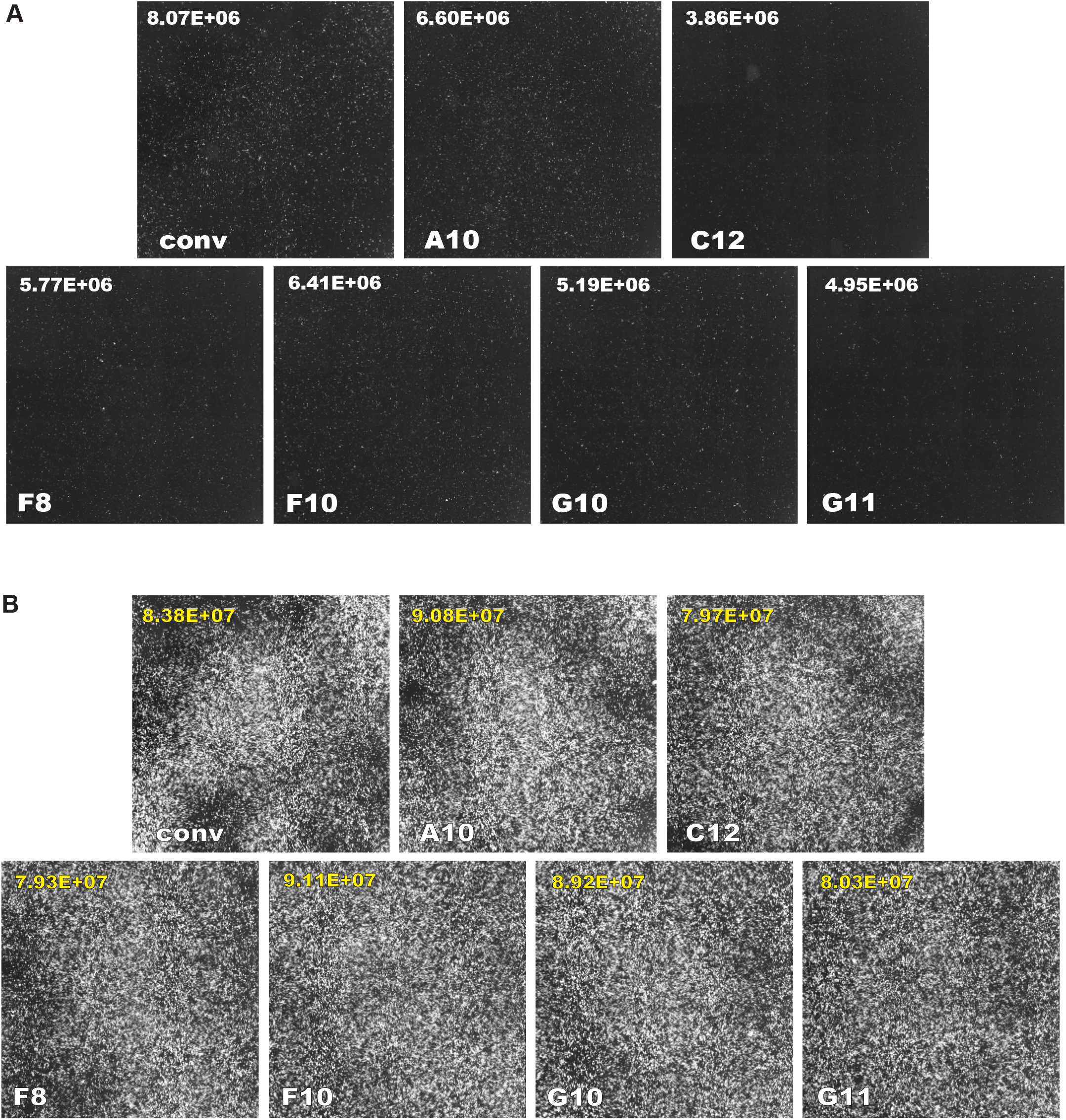
Fluorescence images of new ecDHFR DDs fused to YFP and transiently overexpressed in cells. Images and total fluorescence values from transfected HEK-293A cells 24 h after transient transfection (A) and 24 h after TMP treatment (48 h total after transfection) with constructs containing the conventional or new (A5, C12, F8, F10, G10, and G11) ecDHFR DDs fused to YFP. Numerical values included with each image represent total fluorescence as quantified by Nexcelom Celigo software. n ≥ 3

To more quantitatively confirm the enhanced potential of these new ecDHFR DDs using an orthogonal method, we used western blotting to measure the abundance of the conventional and new mutant ecDHFR DDs (Fig. 3A, B). Comparison of band intensity values indicated that all six ecDHFR DD mutants had significantly lower basal abundance compared to the conventional ecDHFR DD, with the C12 variant having the lowest basal value (Fig. 3B). Moreover, both the A5 and C12 ecDHFR DD mutants exhibited no significant difference in TMP-stabilized levels compared to the conventional ecDHFR DD, indicating that these variants have a larger dynamic range than the conventional ecDHFR DD (Fig. 3B). We next explored the TMP dose-responsiveness of these two ecDHFR DDs to compare their sensitivity to TMP. Transfected HEK-293A cells were treated with 10 nM – 10 μM TMP (24 h) and imaged (Fig. 3C). Both the A5 and C12 mutants behaved indistinguishably from the conventional ecDHFR DD across all TMP concentrations, indicating a similar degree of sensitivity to stabilizer, yet still clearly having lower basal fluorescence than the conventional ecDHFR DD (Fig. 3C). In all subsequent experiments, we focused solely on C12 because of its culmination of favorable characteristics. Moreover, we verified that the unstable C12 ecDHFR DD was degraded by the proteasome (Fig. S2A) and that TMP stabilization occurred post-translationally (Fig. S2B), behaviors which are identical to the conventional ecDHFR DD^4^. Additionally, we also demonstrated that like the conventional ecDHFR DD, C12 is also functional *in vivo* by injecting mouse eyes with AAV containing a C12 ecDHFR DD YFP construct, observing fluorescence in the retina only after administration of TMP (Fig. S3).

**Figure 3.**
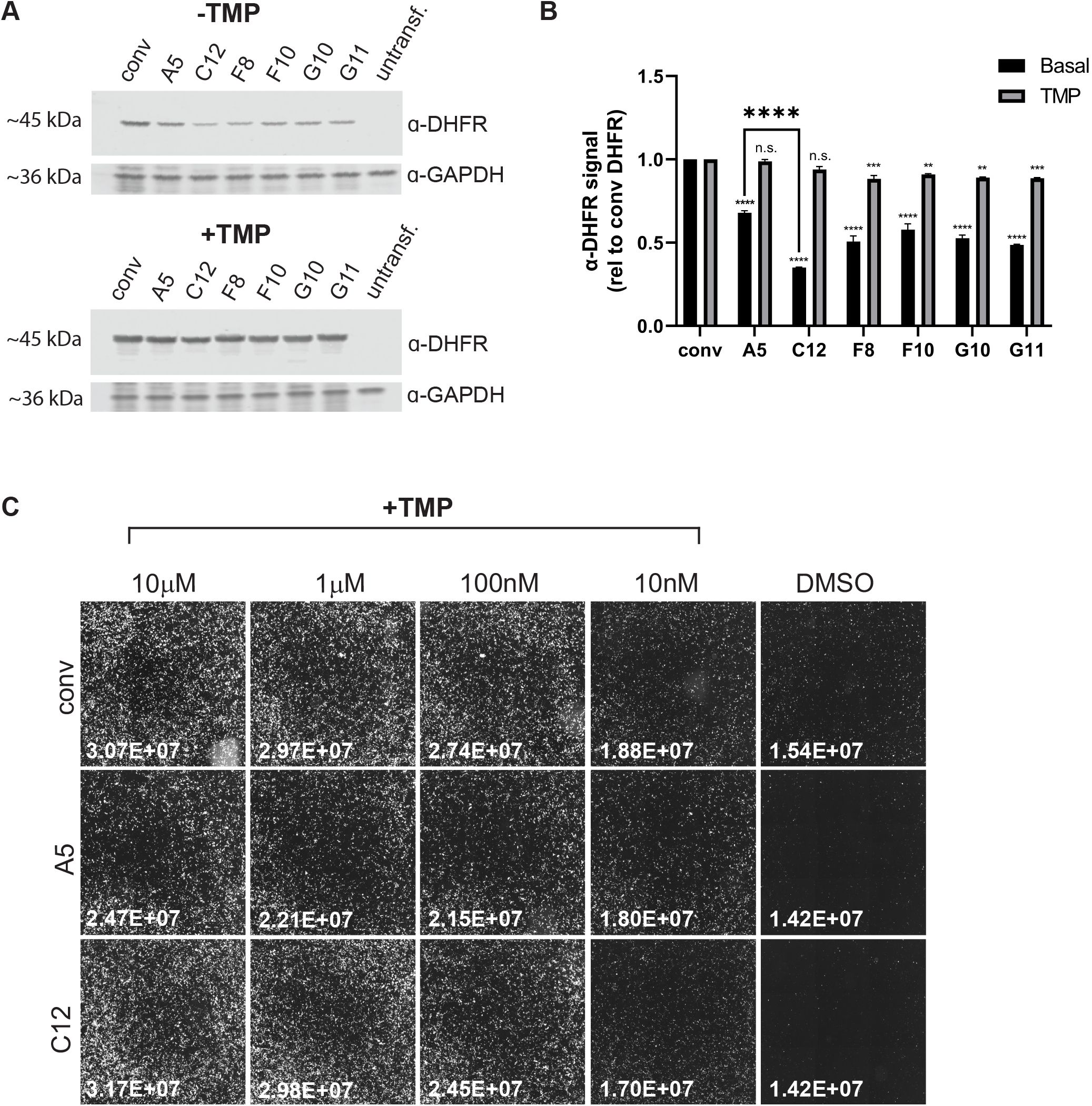
C12 ecDHFR DD exhibits the lowest basal levels and highest TMP sensitivity in transfected cells. (A) Western blot of ecDHFR DDs (conventional, A5, C12, F8, F10, G10 and G11) in transfected HEK-293A cells with and without TMP treatment. Blots were probed with rabbit anti-DHFR and mouse anti-GAPDH. (B) Quantification of band intensities of western blot, with values normalized to that of conventional ecDHFR DD. n = 3, mean ± SEM (**p < 0.01, ***p < 0.001, ****p<0.0001), two-way ANOVA versus conventional ecDHFR samples. (C) Fluorescence images of cells transfected with ecDHFR DDs (conventional, A5 and C12) fused to YFP and treated with decreasing doses of TMP (10 μM, 1 μM, 100 nM, 10 nM, and DMSO neg ctrl) for 24 h. Numerical values included with each image represent total fluorescence as quantified by Nexcelom Celigo software. n = 2

Computational modeling of the C12 ecDHFR DD identifies key residues in promoting destabilization. To explore possible mechanisms for the improved performance of C12 over the conventional ecDHFR DD previously described by Iwamoto *et. al*., we looked at the constructs from a structural perspective. Except for T113S, the C12 mutations are all located distal to ecDHFR’s ligand binding sites (Fig. 4A). Meanwhile, the conventional ecDHFR DD construct features the G67S and Y100I mutation proximal to a ligand binding site, R12Y distal to ligand binding sites (Fig. 4B). To predict the effect of mutations on the stability of ecDHFR both in complex with TMP and NADPH (*holo*-ecDHFR) and without ligands bound (*apo*-ecDHFR), we conducted *in silico* mutagenesis and ddG calculations using Rosetta. We generated 100 minimized wildtype and mutant structures of *holo*- and *apo*-ecDHFR and used the lowest energy of each to estimate the change in ddG between the wildtype and mutant proteins. Inspection of the lowest energy *holo*-wildtype (Fig. 4C, top) and *holo*-mutant structures (Fig. 4C, bottom) for the W74R mutant and T113S mutants (Fig. 4C, left and right panels, respectively) shows that the Rosetta protocol maintains the wildtype protein and ligand conformation accurately (Fig. 4C, left panels) with rmsd’s 0.44-0.45 Å to the native conformation, while the W74R and T113S mutant structures only deviate 0.46 Å from the input structure (Fig. 4C, right panels). The *apo* forms of wildtype and mutant proteins deviated with similar magnitudes (rmsd <0.5 Å, data not shown).

**Figure 4.**
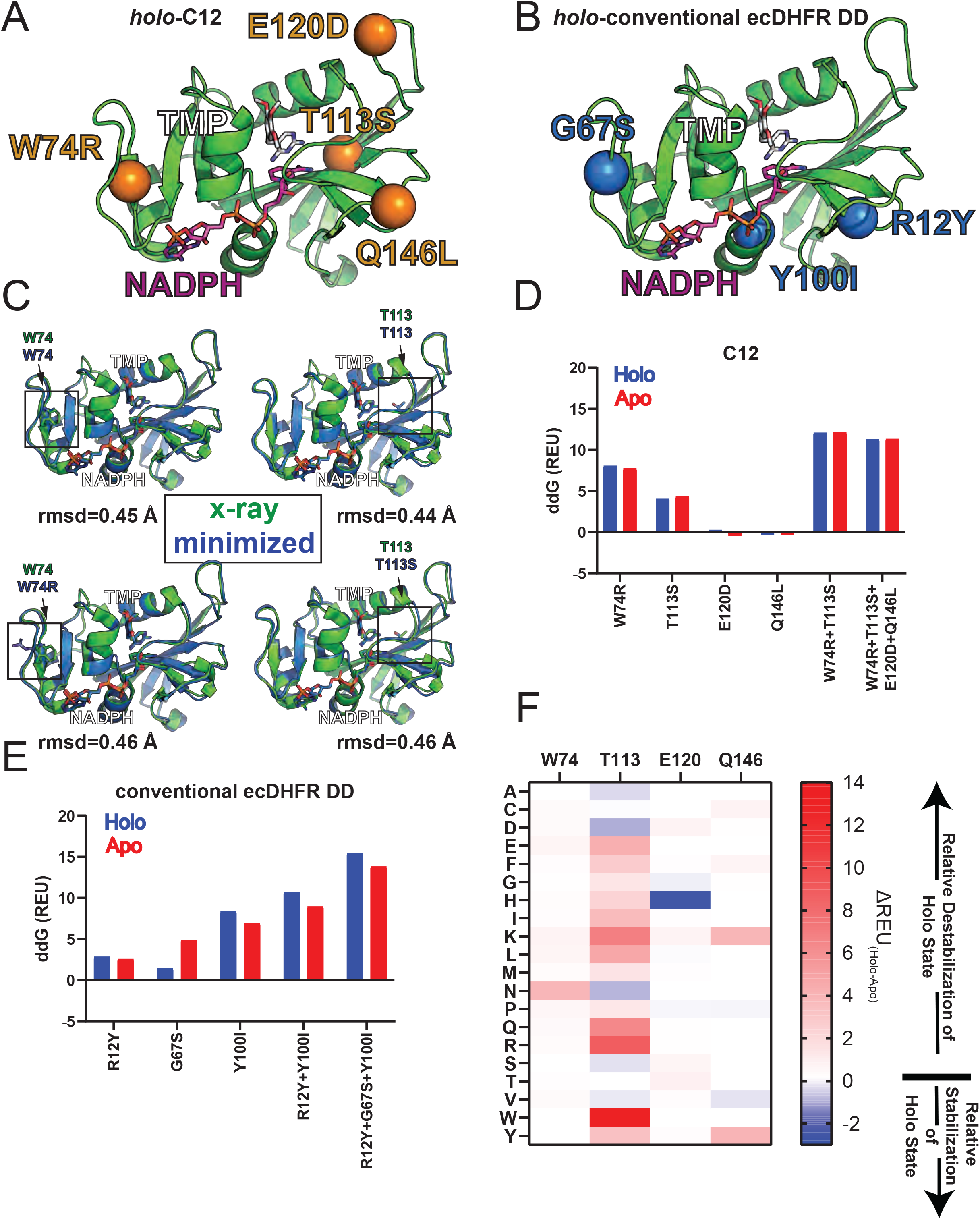
Predicted impact of mutation on ecDHFR stability via *in silico* ddG calculation. (A, B) Illustration of the locations of ecDHFR destabilizing domain mutants on the crystal structure of ecDHFR shown in green (PDB ID: 6XG5), with ligands TMP (white) and NADPH (pink). Locations of C12 mutants are shown in orange (A), while locations of the mutants in the conventional N-terminal ecDHFR DD are shown in blue (B). Sites of mutation are shown as Cα spheres. (C) Superimposition onto the ecDHFR x-ray structure (green) of the minimized wildtype (blue, top half) and mutant (blue, bottom half) structure generated by the RosettaScripts protocol used to calculate ddG of mutation for both the W74 mutant (left half) and the T113S mutant (right half). All-atom rmsds were calculated in pymol. (D,E) Bar plots depicting the impact of mutation(s) on the ΔG of folding as predicted by the Rosetta software suite, represented as ddG_(mutant-wildtype)_ in Rosetta Energy Units (REU). Positive ddG indicates that the mutation is predicted to destabilize the structure compared to the wildtype structure, while negative ddG indicates a predicted increase in stability due to mutation. ddG of mutation are calculated for both the *holo*-ecDHFR complex, bound to both TMP and NADPH (blue), and the *apo*-ecDHFR structure without ligands (red). Predicted impact of mutations is shown for the individual mutations composing C12, the double mutant W74R+T113S, and the full C12 construct (D), as well as the mutations composing the conventional ecDHFR individually, as an R12Y+Y100I double mutant, and the triple mutant R12Y+G67S+Y100I (E). (F) Heatmap depicting the predicted impacts of mutations to C12 mutation sites on the stability of the *holo*-ecDHFR complex relative to *apo*-ecDHFR. The wildtype amino acid and position are shown on the top, while the mutant amino acid is shown on the left. Scale goes from red (*holo*-DHFR is more destabilized by mutation or less stabilized by mutation than *apo*-ecDHFR) to blue *(holo-DHFR* is more stabilized or less destabilized by mutation than *apo*-ecDHFR). White or less saturated colors indicate that the impact of mutation on predicted structural stability is the same or similar for both *holo*- and *apo*-ecDHFR.

This method thus allows us to probe the native state structural energetics of the system and how they are perturbed by mutation.

In general, we find that the C12 mutations destabilize both *holo*- and *apo*-ecDHFR evenly, with most of the destabilizing effect originating from the W74R and T113S mutations (Fig. 4D). This effect is noted both individually, and when both mutations are made concurrently with or without the E120D/Q146L mutations. The E120D and Q146L mutations are predicted to have minimal impact on structural stability. T113S, despite its proximity to the TMP binding site, has minimal differential impact on ecDHFR bound to TMP and NADPH (*holo*-ecDHFR) vs. unbound to its ligands (*apo*-ecDHFR), possibly due to maintenance of the sidechain hydroxyl. Surprisingly, the conventional ecDHFR DD, on the other hand, is predicted to be more destabilized than the C12 construct. The Y100I mutation is predicted to be the most impactful mutation in this construct, and likely due to its proximity to the NADPH binding site, impacts *holo*-ecDHFR energetics more than *apo*-ecDHFR (Fig. 4E). The R12Y and G67S mutations also have smaller contributions to the destabilization of ecDHFR. Interestingly, on its own, the G67S mutation seems to destabilize the unbound state of ecDHFR, but the effect is predicted to be lost when the mutation is present in conjunction with R12Y and Y100I (Fig. 4E). Additionally, the conventional ecDHFR DD construct was also more destabilized relative to wildtype when bound to ligands than when not bound to ligands, indicating a narrowing of the stability gap between the *holo*-ecDHFR and *apo*-ecDHFR states.

To further probe the C12 and conventional construct mutation sites, we calculated the ddG of mutations made to these sites individually using *in-silico* saturation mutagenesis (Fig. S4). We find that most substitutions at W74, E120, Q146, and R12 (Fig. S4A, C, D, and F respectively) destabilize the protein as a whole and do not differentially destabilize ecDHFR bound to its ligands respective to unbound ecDHFR. Mutations at positions T113 and Y100 generally destabilize *holo*-ecDHFR relative to *apo*-ecDHFR, as might be expected from their locations proximal to ligand binding sites (Fig. S4B and G). Mutation in Rosetta to amino acids with similar properties to the native sequence have negligible effects on the stability, for example, aromatic substitutions at position 74 (Fig. S4A), or substitutions to threonine or valine at position 113 (Fig. S4B) have no effect on stability. Interestingly, G67, located on a flexible loop, possesses several mutants that destabilize the unbound complex more than the bound complex, possibly due to sidechain interactions with NADPH phosphates compensating for the loss in loop stability (Fig. S4E). When we directly compare the ddG of mutations between *holo*- and *apo*-ecDHFR, we see there is minimal differential destabilization in mutations to W74, E120, and Q146, which are residues with minimal ligand interactions. However, mutations to T113, a residue involved in TMP binding, changes the stability gap between *apo*- and *holo*-ecDHFR (Fig. 4F).

### The C12 variant requires all four missense mutations for its enhanced basal degradation and/or full TMP stabilization

To better understand the role of each missense mutation that comprises the C12 variant, in parallel wet-lab experiments, we used site-directed mutagenesis to revert each individual missense mutation to their wildtype ecDHFR counterpart residue (i.e., R74W, S113T, D120E, L146Q). Fluorescence images and quantification of cells transfected with these plasmids revealed that the reversal of the R74, S113, and D120 mutations elevate basal YFP fluorescence, and are thus likely combinatorially responsible for promoting ecDHFR instability and/or proteasomal targeting (Fig. 5A, B). Cells expressing the C12 L146Q reversal mutation have similar basal YFP signal compared to the unmodified C12 ecDHFR DD (Fig. 5A), but have a lower induced YFP signal (Fig. 5B), indicating that this residue is not particularly important for basal turnover, but rather for inducibility. The R74W, S113T and D120E reversions also appear to limit the extent of TMP-induced stabilization, but to a lesser degree compared to L146Q (Fig. 5B). Overall, these data suggest that the four missense mutations in C12 work in concert with each other—some more important than others—the combination of which promotes basal degradation and TMP inducibility.

**Figure 5.**
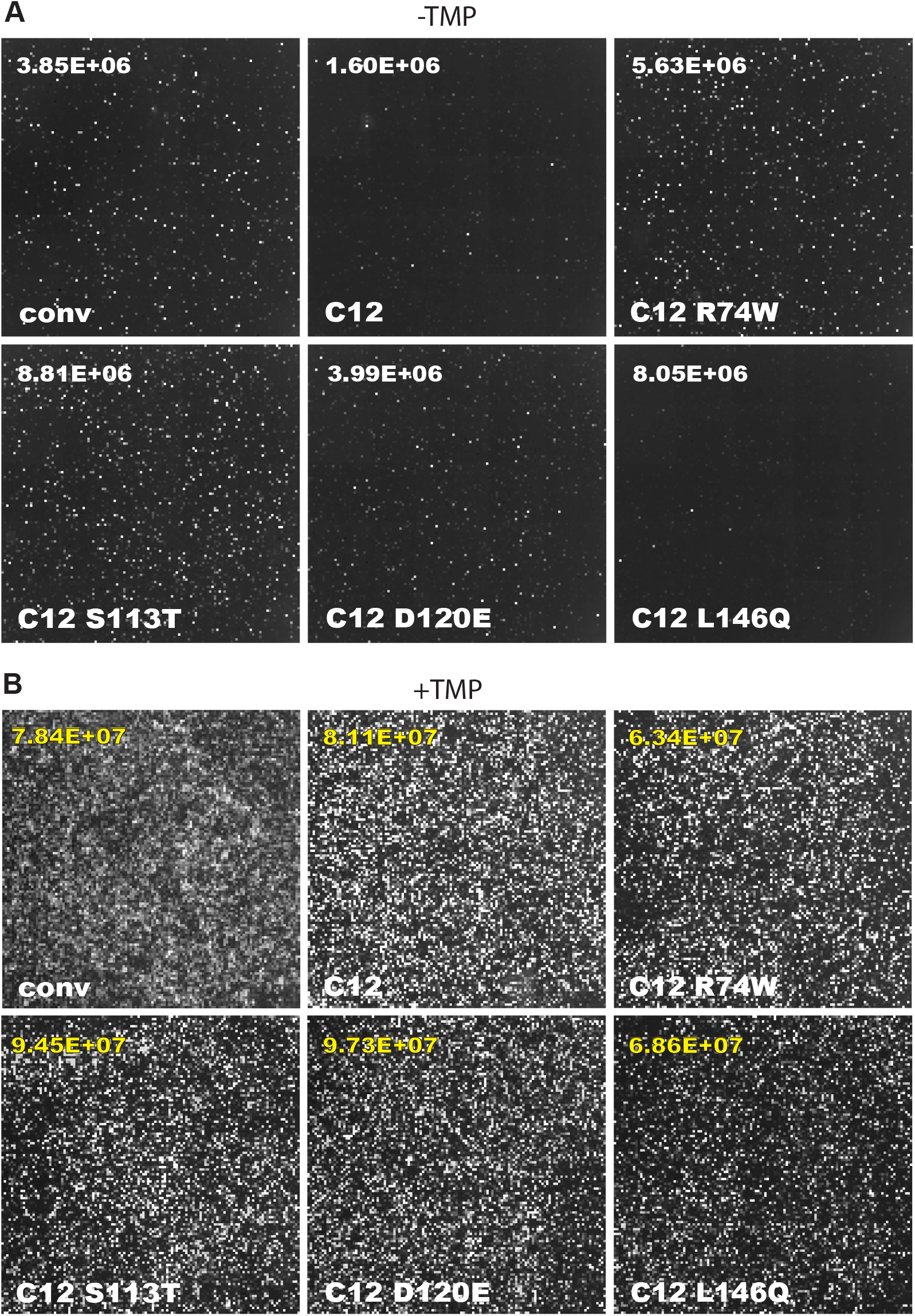
Individual mutations of C12 ecDHFR DD each contribute to its DD capabilities. Fluorescence images and quantitation of HEK-293A cells transfected with individual mutations of C12 ecDHFR DD before (A) the addition of TMP (B). Numerical values included with each image represent total fluorescence as quantified by Nexcelom Celigo software. n = 3

### C12 requires the full length ecDHFR sequence to achieve both high levels of basal turnover and TMP-inducibility

We next determined whether we could truncate the C12 ecDHFR DD and still retain its enhanced basal degradation and TMP stabilization features. Generating a smaller drug-controllable DD is an attractive goal, especially when packaging genetic information into AAV-based gene therapies with a ~5 kb size limit^18, 19^. To accomplish this goal, we developed a series of truncation variants that harbored C12 as the core degron, but lacked either N-terminal or C-terminal regions. We generated plasmids with various truncated versions of the C12 ecDHFR DD (M1-D120, S64-P130, G67-R159, R74-S113, R74-D120) fused to YFP and transfected them into cells (Fig. S5A, B). Fluorescence images and quantifications of these cells before and after TMP treatment illustrate that despite the inclusion of important mutations, each truncation falls short of meeting the DD qualities of the unmodified C12 ecDHFR, either in destabilization or TMP-induced stabilization. While the M1-D120 and G67-R159 mutants have lost the latter feature, the S64-P130, R74-S113 and R74-D120 mutants have lost the former (Fig. S5A, B). These results emphasize the difficulty in achieving a substantial shortening to the ecDHFR DD sequence without compromising its most important regulatory features.

### Controlling NF-κB and Nrf2 signaling using the C12 ecDHFR DD

Next we tested whether C12 could be confer its enhanced degradation abilities to more biologically relevant proteins in addition to YFP, translating into improved control over downstream gene activation as well. The two proteins we used to test this behavior were IκBα and Nrf2, both of which are involved in carefully regulated cellular pathways and target inflammation and oxidation (two phenomena closely linked to retinal degeneration^20, 21^), respectively. We generated constructs expressing either conventional or C12 ecDHFR DD fused at the N terminus of either IκBα or Nrf2ΔNeh2/Neh6, a truncated but biologically functional version of Nrf2 that lacks the Neh2 and Neh6 negative regulatory domains, allowing for enhanced activation^22, 23^. Cells transfected with these constructs were treated with either TMP or DMSO control (24 h), then analyzed for western blotting to quantify for IκBα or Nrf2 (Fig. 6A-C and Fig. 7A-C). In both cases, C12 conferred better destabilization and therefore significantly lower basal levels of IκBα and Nrf2, in comparison to the conventional ecDHFR DD, while still maintaining similar levels of TMP-induced stabilization (Fig. 6A-C and Fig. 7A-C). Overall, C12 commanded a wider dynamic range in controlling IκBα and Nrf2 than their conventional counterparts (Fig. 6C and 7C). Moreover, C12’s enhanced degradation ability was also manifested in assessing levels of heme oxygenase-1 (HO-1), a protein whose transcription is strongly regulated by Nrf2 (Fig. 7A, D-E), in agreement with a similar trend as previous experiments monitoring the client proteins, IκBα and Nrf2. Overall, these results demonstrate the improved control over various proteins and their downstream partners using C12 versus the conventional N-terminal ecDHFR DD.

**Figure 6.**
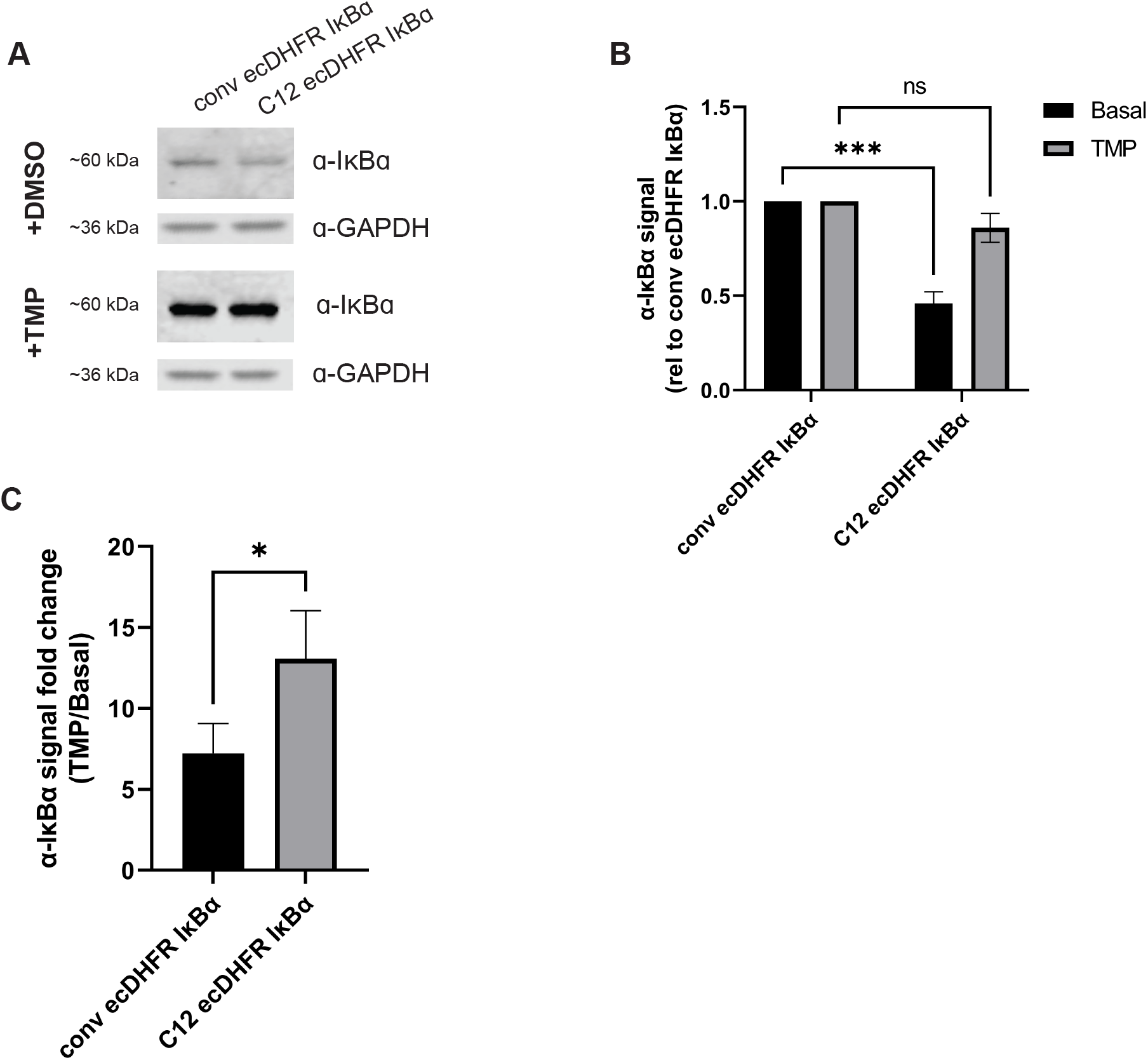
C12 ecDHFR DD control of IκBα demonstrates enhanced basal degradation. (A) Western blot of DMSO versus TMP treated cells transfected with the conventional ecDHFR DD or C12 fused to IκBα. Blots were probed with rabbit anti-IκBα and mouse anti-GAPDH. (B) Band intensities were quantified (via LI-COR Image Quant software) relative to the conventional ecDHFR DD samples. (C) Fold change was calculated by dividing band intensity values of TMP-treated samples with the corresponding value from DMSO-treated samples. n ≥ 3, mean ± SEM (*p < 0.05, **p < 0.01, ***p < 0.001), two-way ANOVA versus conventional ecDHFR samples (B) and RM one-way ANOVA for (C).

**Figure 7.**
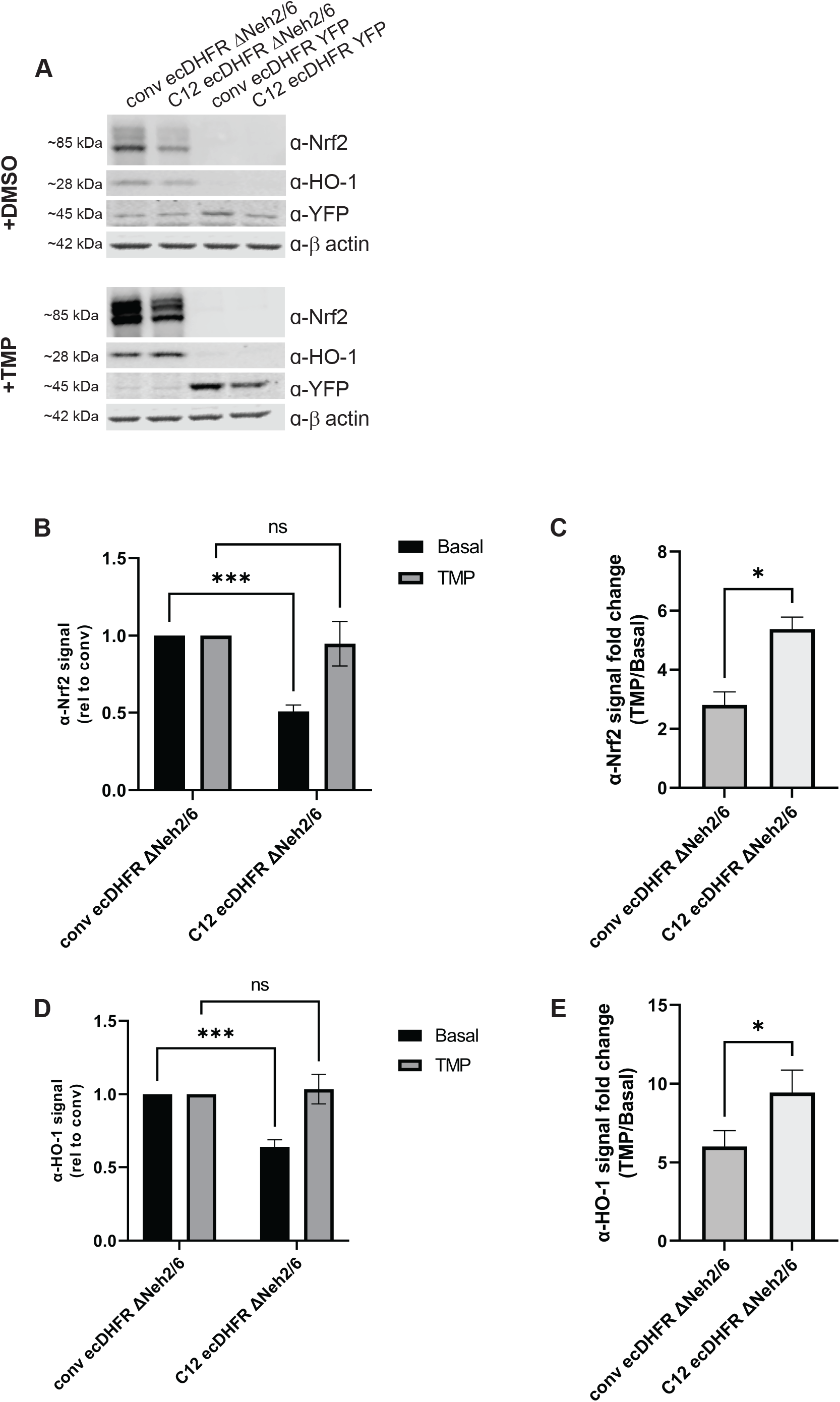
C12 ecDHFR DD fusion to Nrf2 demonstrates enhanced control over fusion protein levels and downstream HO-1. (A) Western blot of DMSO versus TMP treated HEK-293A cells transfected with conventional ecDHFR DD or C12 Nrf2△Neh2/6, and conventional ecDHFR DD or C12 YFP. Blot was probed with rabbit anti-Nrf2, mouse anti-HO-1, mouse-anti GFP, and rabbit anti-β actin. Band intensities were quantified (via LI-COR Image Quant software) relative to the conventional ecDHFR DD samples for Nrf2 (B) and HO-1 (D). Fold change was calculated through dividing band intensity values of TMP-treated samples with the corresponding value from DMSO-treated samples for Nrf2 (C) and HO-1 (E). n ≥ 3, mean ± SEM (*p < 0.05, **p < 0.01, ***p < 0.001), two-way ANOVA versus conventional ecDHFR samples (B and D) and RM one-way ANOVA for (C and E).

## DISCUSSION

Our work builds upon improving the ecDHFR DD system by developing and screening through a library of ecDHFR mutants and identifying a new ecDHFR DD, dubbed C12, with enhanced destabilization capabilities and analogous TMP-induced stabilization properties to the conventional ecDHFR DD. We followed a similar approach to Iwamoto *et al’s* development of the original ecDHFR DD^4^, using error-prone PCR with an E. coli DHFR template, and fusing the library to a YFP reporter for screening of stabilization/destabilization (Fig. 1). The diversity in mutations that allow for ecDHFR’s DD-like behaviors suggest the considerable tolerance of its TMP binding ability to mutations. However, our truncations to the ecDHFR DD domain do severely compromise its essential features, either its inherent instability or TMP binding (Fig. S5A, B). Previous work has developed a truncated version of the conventional ecDHFR DD to serve as a degron in AAV-based gene therapy, in which the last 54 amino acids (which include those involved in TMP binding) were removed, leaving an M1-P105 ecDHFR DD degron^18^. In a similar vein, the M1-D120 and G67-R159 C12 ecDHFR truncations (the latter of which is smaller than the M1-P105 truncation) have the potential to serve as a degron (Fig. S5A, B), which could be useful in the context of the ~5kb size limit in AAV for gene therapy.

Furthermore, our success in identifying a list of unique TMP-responsive ecDHFR mutants suggests the versatility of the library screening process for discovering new DDs and stabilizers. Previously, we have screened a library of compounds in cells stably expressing the conventional ecDHFR DD, and discovered several effective alternative stabilizers to TMP^12^. A similar process can be applied for screening new destabilizing domains responsive to a particular stabilizer of interest (e.g., a TMP analog or similar structure^24^). For example, the original dark population obtained as described in Fig. 1 expresses a variety of destabilized ecDHFR mutants that could be further screened by treatment with said stabilizer, followed by another round of FACS to isolate fluorescent clones, thus allowing for the discovery of new DDs responsive to select stabilizers. The random mutagenesis itself may also be performed on other protein domains with known binding partners, facilitating the discovery of new DDs, as was the case in the development of the human estrogen receptor ligand binding domain (ERLBD)-based DD^5^. Moreover, the consistency between *in silico* versus experimental results in determining the importance of mutations for ecDHFR’s stability opens the possibility of using computational models to predict hot spot sites for destabilization selective for unbound ecDHFR, thus informing how to generate the ‘ideal’ DD.

One major question that arises with the comparison between the C12 and conventional ecDHFR DD is the biochemical explanation for why C12 overall behaves better than the conventional ecDHFR DD. Ideally, a reversible DD should have a sufficiently high-energy, unstable structure when out of complex with its regulatory ligand in order to promote unfolding and subsequent recognition/degradation by the cellular proteasome, while being sufficiently stable when bound to its ligand to reduce/avoid degradation. When engineering destabilizing domains, these factors also have to be balanced with maintaining ligand binding affinity and optimizing protein degradation rates in the unbound state to have sufficiently short half-life while maintaining structure in the ligand bound state. This may form the biophysical basis for C12’s performance relative to the Iwamoto *et. al*. construct - although it is not as destabilized as the previously described ecDHFR DD, the stability gap between its ligand-bound and unbound states is higher, possibly leading to more targeted, differential degradation of TMP-unbound C12 ecDHFR and sustained stability of TMP-bound C12 ecDHFR. Alternatively, additional unaccounted for biochemical properties could explain C12’s improved control of fused proteins. For example, the C12 ecDHFR may be polyubiquitinated more easily than the conventional ecDHFR DD, or perhaps be targeted by additional ubiquitin-independent proteasome degradation pathways. Future co-immunoprecipitation experiments aimed at isolating the C12 ecDHFR DD and its binding partners could help elucidate the nature of its enhanced degradation, or even conserved features that allow for the DDs to be better degraded in the first place.

An important aspect of the C12 ecDHFR DD is that its enhanced degradation feature remains consistent when fused to YFP, IκBα, or Nrf2, and is reflected in the downstream partners of Nrf2 as well (Fig. 6A-C and Fig. 7A-E), thus emphasizing the functional benefit of using the C12 over the conventional ecDHFR DD. Previously, we have used the conventional ecDHFR to control IκBα in order to suppress NFκB activation in human immortalized RPE cells; however, detectable basal stabilization of the ecDHFR IκBα fusion protein led to the need for an additional doxycycline-inducible control measure for the fusion protein^14^. A similar issue of basal stabilization occurs when the conventional ecDHFR is fused to Nrf2 (unpublished results). In such cases, the C12 ecDHFR DD can serve as an effective replacement to potentially solve these issues. With the successful use of the conventional ecDHFR DD in past applications, the C12 ecDHFR DD can only further increase the opportunities for the system’s utilization.

Still, there remain gaps in verifying the C12 ecDHFR DD’s true functional applicability within the context of gene therapy and intervention of complex biological pathways with multiple regulatory elements. One limitation of this paper is in the lack of deeper investigation into therapeutic benefit from the application of the C12 ecDHFR DD in more relevant cell culture systems. Follow up studies should explore whether the control of IκBα or Nrf2 using C12 ecDHFR DD could lead to substantial improvement in anti-inflammatory regulation or oxidative stress relief, respectively, in model systems besides HEK293A cells. For example, an interesting prospect to explore is whether using the C12 ecDHFR DD to inducibly activate Nrf2 could lead to the improved oxidative stress relief over the constitutively expressed Nrf2, which has been observed in the doxycycline-inducible version in retinal pigmented epithelial cells^25^.

Overall, our work in developing and verifying the C12 ecDHFR DD marks an important milestone highlighting a second-generation ecDHFR DD system. We are hopeful that the improvements made to this DD will encourage wider use of the system and will enable its translational utility in gene therapy applications as well.

## MATERIALS AND METHODS

### Library Generation and Screening

An E. coli dihydrofolate reductase (ecDHFR) random mutagenesis library was generated by error-prone PCR (~8-10 mutations per kb) from the wildtype ecDHFR cDNA template (GenScript, Piscataway, NJ). The library was cloned into a pENTR1A plasmid backbone with a C-terminal YFP HA reporter tag using Gibson Assembly (HiFi DNA Assembly Cloning Kit, New England Biolabs, (NEB), Ipswich, MA), generating ~1200 individual colonies. Individual colonies were collected and combined, and then shuttled as a mixture of DNAs into the pLenti CMV Puro DEST vector (gift from Eric Campeau and Paul Kaufman, Addgene plasmid # 17452) via an LR clonase II reaction (Invitrogen, Waltham, MA). VSV-G-pseudotyped lentivirus was generated by cotransfection of HEK-293T cells (CRL-3216, ATCC, Manassas, VA) with psPAX2 (gift from Didier Trono, Addgene plasmid #12260), pMD2.G (gift from Didier Trono, Addgene plasmid #12259), and the pLenti library as described previously^26^. To generate the ecDHFR-expressing cell library, Chinese hamster ovary cells (CHO/dhFr-, CRL-9096, ATCC) were seeded into a 10 cm dish (Costar, Corning, Corning, NY) at a density of 3 million cells/dish and allowed to attached overnight. The following day, cells were replenished with fresh media containing 1 μg/mL polybrene and 250 μL of crude lentivirus, and left to incubate for 24 h to achieve a multiplicity of infection (MOI) of ~1. Infected cells were then selected with puromycin (10μg/mL) for 2 weeks.

CHO cells stably expressing the ecDHFR mutant library were submitted for flow cytometry twice (FACSAria, (BD Biosciences, Franklin Lakes, NJ), UT Southwestern Mass and Flow Cytometry Core) to isolate a YFP-negative population of cells lacking fluorescence. This population was then treated with TMP (10 μM) and sorted 24 h later for YFP-positive cells with fluorescence into black clear-bottomed 96 well plates (Nunc, Roskilde, Denmark), with a single cell per well. Once single celled clones had grown into confluent colonies, lack of YFP fluorescence in the absence of stabilizer was verified by imaging wells on the Celigo Imaging Cytometer (Nexcelom Biosciences, Lawrence, MA) for YFP fluorescence. Next, all wells were treated with TMP for 24 h and imaged on the Celigo again to confirm which clones exhibited strong YFP fluorescence. Clones with the best response to TMP were expanded and lysed for gDNA extraction and purification using the QuickExtract DNA extraction solution (Lucigen, Middleton, WI). Genomic ecDHFR was amplified (forward: CCAAGTACGCCCCCTATTGA reverse: ATATCTCGAGTGCGGCCGCGTTATGCGTAGTCTGGTACGTCG) and purified from the gDNA template using Q5 High-Fidelity DNA Polymerase (NEB) and QIAquick PCR purification kit (Qiagen, Hilden, Germany), respectively, for subsequent analysis and cloning.

### Plasmid Cloning and Mutagenesis

New ecDHFR DDs (clones A5, C12, E8, F8, F10, and G11) were shuttled into a pcDNA vector with a YFP reporter via Gibson Assembly. Single and multiple point mutation modifications to the pcDNA C12 construct were generated through the Q5 Site-directed Mutagenesis kit (NEB) or purchased as a minigene from Integrated DNA Technologies (IDT, Coralville, IA). Constructs of truncated C12 DHFR and C-terminal C12 DHFR were also purchased as minigenes (IDT). All minigene constructs were subsequently cloned into pcDNA through restriction digest (XmnI and SphI (NEB)) and ligation into a pcDNA vector. The C12 DHFR IκBα construct was generated through restriction digest (SphI and XbaI (NEB)) and ligation of C12 ecDHFR into a digested pENTR1A IκBα backbone described previously^14, 20^. The C12 DHFR FLAG-tagged (FT) Nrf2 lacking the Neh2/Neh6 regulatory domains (ΔNeh2/6) construct was generated via Gibson Assembly. AAV constructs (C12 and conventional ecDHFR^4^ YFP 2A FLuc) were generated by Gibson Assembly into a pENTR1A backbone, and later shuttled to a pAAV Gateway vector (gift of Matthew Nolan, Addgene plasmid #32671) through LR recombination (Invitrogen). All constructs assembled via Gibson Assembly (i.e., HiFi DNA Assembly, NEB) were designed using the NEBuilder online software (https://nebuilder.neb.com/). All constructs were fully verified by Sanger sequencing, analyzed with SnapGene 5.1.7 (GSL Biotech, San Diego, CA, USA), and purified with the Qiagen Plasmid Plus Midiprep kit (Qiagen) for transfections.

### Cell Culture and Transfections

HEK-293A cells (R70507, Life Technologies, Carlsbad, CA) and HEK-293T cells (ATCC) were cultured in Dulbecco’s high-glucose minimal essential medium (DMEM, 4.5 g/L glucose, Gibco, Waltham, MA) supplemented with 10% fetal bovine serum (FBS, Omega Scientific, Tarzana, CA) and penicillin-streptomycin-glutamine (PSQ, Gibco). CHO cells were cultured in Iscove’s Modified Dulbecco’s Medium (IMDM, ATCC) supplemented with 10% FBS, PSQ and hypoxanthine-thymidine (ATCC). Cells were grown at 37°C and 5% CO_2_.

For transient transfections, HEK-293A cells were seeded in 12, 24 or 48 well plates (Costar, Corning) at a concentration of 110,000, 75,000 and 35,000 cells per well, respectively. DNA plasmids were introduced into cells using Lipofectamine 3000 (Life Technologies, Carlsbad, CA), at a concentration of 1 μg/well (3 μL of L-3000 and 2 μL P3000) in 12-well plates, 500 ng/well (1.5 μL L-3000 and 1 μL P3000) in 24 well plates, and 250 ng/well (0.75 μL L-3000 and 0.5 μL P3000) in 48 well plates..

### Fluorescence Imaging for Transient Transfections

To decrease background autofluorescence signal due to phenol red, cell culture media was changed to Fluorobrite DMEM (Gibco) supplemented with 1% FBS and PSQ 24 h after transfection. Cells were then imaged for baseline YFP fluorescence at 150000 ms exposure on the Celigo Nexcelom. Cells were treated with 10 μM TMP and imaged again under the same parameters for stabilized YFP fluorescence 24 h later.

### Western Blotting

Transfected cells were washed with HBSS and lysed in-well with radioimmunoprecipitation assay (RIPA) buffer (Santa Cruz, Dallas, TX) supplemented with Halt Protease Inhibitor (Pierce, Rockford, IL) and benzonase (Millipore Sigma, St. Louis, MO, USA) for 15 minutes at RT. Lysate was then centrifuged at 14,000 rpm for 10 min at 4°C, followed by collection of the soluble fraction for analysis of protein concentration via bicinchoninic assay (BCA, Pierce). Fifteen to 20 μg of protein was loaded into a 4-20% Tris-Gly SDS-PAGE gel (Life Technologies) and transferred onto a nitrocellulose membrane with the iBlot 2 semidry transfer system (Life Technologies). Membranes were stained with Ponceau S (Sigma) to assess for total protein transfer, followed by overnight blocking in Odyssey Blocking Buffer (LI-COR, Lincoln, NE). Western blots were all imaged on the Odyssey CLx (LI-COR) and analyzed with ImageStudio Software (LI-COR).

### In silico mutagenesis and ddG calculation

Rosetta (vers. 3.12) was used to perform structure optimization, mutagenesis, and ddG calculation. The DHFR structure was obtained from the PDB (PDB ID: 6XG5) and optimized for further use in Rosetta using the Rosetta relax application with flags “-in:file:fullatom -ex1 -ex2 -use_input_sc -flip_HNQ -no_optH false -relax:fast -score:weights ref2015”^27^. 500 relaxed structures were obtained, and the lowest energy structure used for subsequent mutagenesis and ddG calculation. The ddG of mutation was calculated using RosettaScripts^28^. The protocol used was derived from the methodology of the Rosetta ddG monomer application, but with fewer restraints^29^. Operating on the preminimized structure, the protocol involved generating both 100 wildtype structures, subject to localized repacking within 20 Angstroms of mutation sites followed by unrestricted global minimization, and 100 mutant structures, in which the desired mutation(s) was introduced prior to localized repacking and unrestricted global minimization. Initial preminimization, repacking, minimization, and scoring were all conducted using the Rosetta Ref2015 scorefunction^30^. The ddG of mutation for a given mutant was calculated by taking the lowest energy wildtype and mutant structures out of the 100 replicates of each as follows:

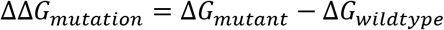

This procedure was repeated for an *apo-state* version of DHFR by taking the preminimized DHFR structure and removing the NADPH and TMP ligands prior to *in silico* mutagenesis and ddG calculation. Afterwards, the impact of mutation on the stability of the *holo*-DHFR complex relative to the *apo*-DHFR structure was calculated using:

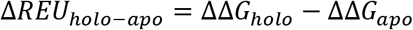

### AAV production and purification

To generate AAV, roughly 5 million AAV pro293T cells (Takara, San Jose, CA) in 30 mL of media were seeded overnight in a T175 flask (Greiner Bio-One, Kremsmünster, Austria). Next, a co-transfection mix was prepared with 3 mL of DMEM (Gibco) containing polyethylenimine (PEI) (3 μg per 1 μg of transfected DNA) and the following three plasmids in a 1:1:1 molar ratio: the AAV construct (C12 or conventional ecDHFR YFP 2A FLuc), pAAV Helper (CellBioLabs, San Diego, CA) vector, and pAGC AAV 2/2 MAX rep cap plasmid^31, 32^. AAV was harvested 96 h post transfection using chloroform extraction and 50% polyethylene glycol (PEG) 8000 (BioUltra, Sigma) precipitation for 1 h, followed by the addition of 1 μL benzonase (25-29 U/L) (Millipore Sigma) and 3.5 1M MgCl_2_. AAV was purified by loading into an iodixanol gradient (15-60%) into QuickSeal Tubes (Seton Scientific, Petaluma, CA), followed by ultracentrifugation on the T1270 rotor at ~280,000xg for 90 min at 4°C (Thermo Scientific). After ultracentrifugation, purified AAV was collected from the 40% iodixanol gradient layer and loaded into the Amicon Ultra 0.5 mL 100K Centrifugal Filter (EMD Millipore, Burlington, MA) for further concentration of AAV, and final elution into HBSS and 0.014% Tween-20 (HBSS-T). To verify purity, 2 μL of purified, concentrated virus was run on a 4-20% Tris-Gly SDS-PAGE gel (Life Technologies), which was then analyzed using the Pierce Silver Stain Kit (Pierce) for visible bands of VP1, VP2, and VP3 from wells of loaded virus. To verify titer, 1 μL of purified virus was analyzed with the Quant-it PicoGreen dsDNA Assay (Invitrogen).

### Intravitreal Injections

Ten-week-old WT C57BL/6 mice were anesthetized with a ketamine/xylazine cocktail (120 mg/kg and 16 mg/kg, respectively) followed by application of tropicamide (dilator, 1%[w/v]) (Alcon, Fort Worth, TX), and GenTeal Severe Dry Eye Gel (Alcon) to maintain ocular hydration. Injections were performed on a stereo microscope (Zeiss, Oberkochen, Germany) using a 33G 1/2” needle attached to a Hamilton micro-syringe (Hamilton, Reno, NV) containing 1 μL AAV (1×10^11^vg/mL), or HBSS-T vehicle control as described previously in greater detail^33^. The eye was proptosed, and punctured above the limbus with a 27G needle, followed by slow injection of the virus into the puncture site at a 45° angle. Following injection, AK-POLY-BAC antibiotic ointment (Akorn, Lake Forest, IL, USA) was applied onto eyes for recovery.

### Fundus Imaging

To measure for ecDHFR YFP fluorescence, intravitreally injected mice were anesthetized with isoflurane gas, followed by application of tropicamide and GenTeal Severe Dry Eye gel for dilation and hydration of the eye. Baseline fundus blue autofluorescence images were first taken on the Spectralis-OCT (Heidelberg Engineering, Heidelberg, Germany) before given TMP in drinking water (400 mg/L). After 48 h with access to TMP water, mice were again imaged for fundus fluorescence images to assess TMP-induced YFP stabilization.

## ACKNOWLEDGEMENTS

JDH is supported by an endowment from the Roger and Dorothy Hirl Research Fund and R01 EY027785. LAJ is the Marie Effie Cain Scholar in Medical Research and is supported by the Chan Zuckerberg Initiative (2018-191983). Additional support was provided by a National Eye Institute Visual Science Core Grant (P30 EY030413, to the UT Southwestern Department of Ophthalmology).

## FIGURE LEGENDS

**Supplemental Figure 1. Sorted clonal populations expressing distinct ecDHFR DD mutants demonstrate a variety in TMP-induced fluorescence intensity.** Fluorescence image of 96 well plate containing clonal populations derived from single sorted cells treated with TMP for 24 h.

**Supplemental Figure 2. Predicted impact of saturation mutagenesis on C12 and conventional ecDHFR DD mutation sites on the stability of complexed and uncomplexed ecDHFR.** (A-F) Bar plots depicting the impact of mutation(s) on the ΔG of folding as predicted by the Rosetta software suite, represented as ddG(mutant-wildtype) in Rosetta Energy Units (REU). Positive ddG indicates that the mutation is predicted to destabilize the structure compared to the wildtype structure, while negative ddG indicates a predicted increase in stability due to mutation. ddG of mutation are calculated for both the *holo*-DHFR complex, bound to both TMP and NADPH (blue), and the *apo*-ecDHFR structure without ligands. *In silico* saturation mutagenesis was carried out and ddG calculated for C12 mutation sites W74 (A), T113 (B), E120 (C), and Q146 (D), as well as the conventional ecDHFR DD sites R12 (E), G67 (F), and Y100 (G).

**Supplemental Figure 3. C12 ecDHFR DD is degraded proteasomally and stabilized cotranslationally.** Fluorescence images and quantitation of cells transfected with C12 YFP and treated with either 10 μM MG132, 100 μM chloroquine (CQ), or no drug (panel A), or with cycloheximide (CHX) or DMSO for 3h followed by TMP for 6 h (panel B). n = 2.

**Supplemental Figure 4. C12 ecDHFR DD is functional *in vivo*.** Infrared (IR) and blue laser autofluorescence (BAF) fundoscopy images of mouse eye captured ≥ 6 weeks after injection of C12 DHFR YFP AAV (n = 3 eyes) or HBSS-T negative control (n = 1), before and after TMP treatment via drinking water (400 mg/L) for 48 h.

**Supplemental Figure 5. Truncations to C12 ecDHFR compromises its integral features.** Fluorescence images and quantitation of HEK-293A cells transfected with various truncations of C12 ecDHFR DD (M1-D120, S64-P130, G67-R159, R74-S113, and R74-D120) before the addition of TMP (A) and after (B, 24 h 10 μM). Numerical values included with each image represent total fluorescence as quantified by Nexcelom Celigo software. n = 3

