## Supplementary figures and images for "Development of a new DHFR-based destabilizing domain with enhanced basal turnover and applicability in mammalian systems"

### supplementary info

Fig. S1

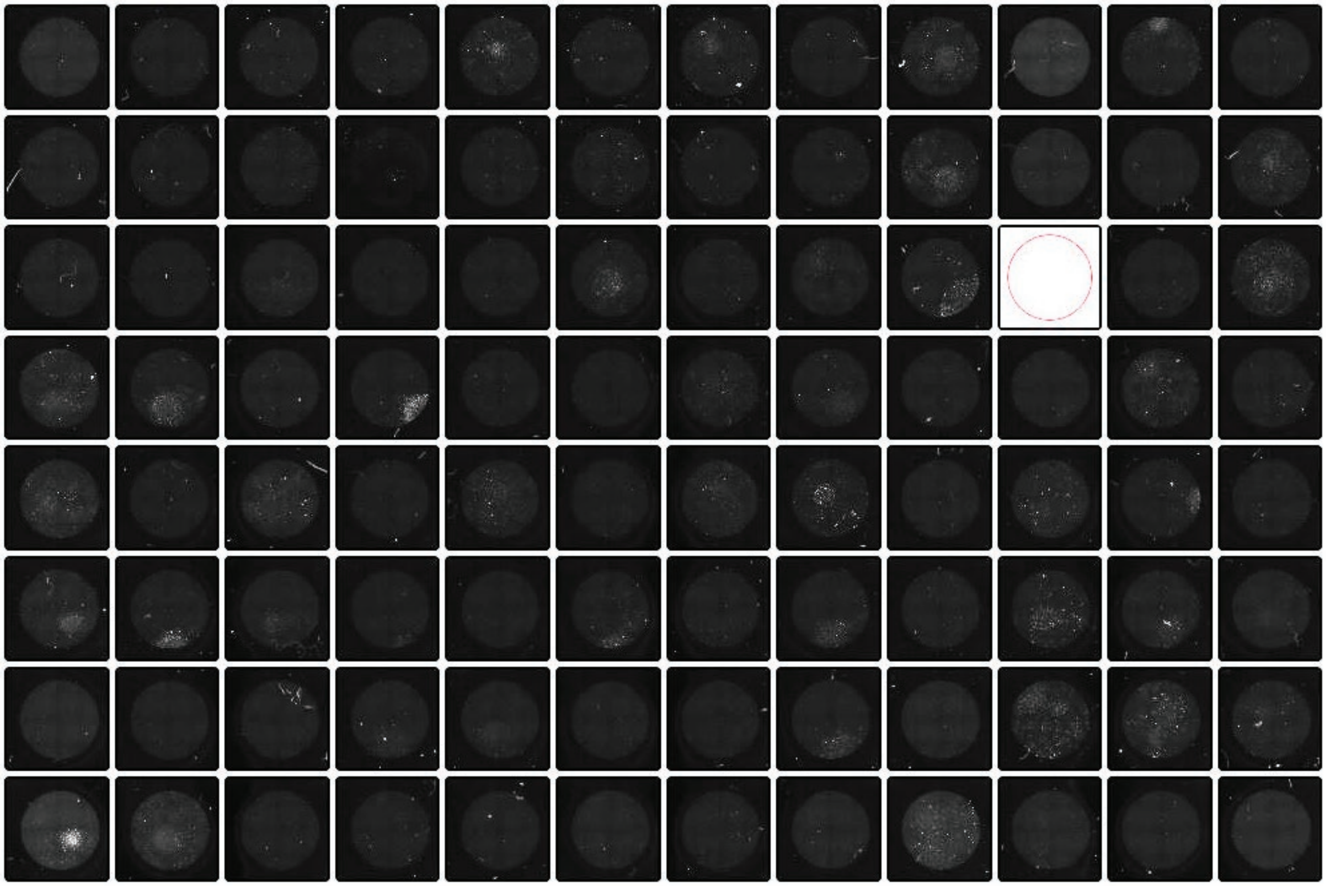

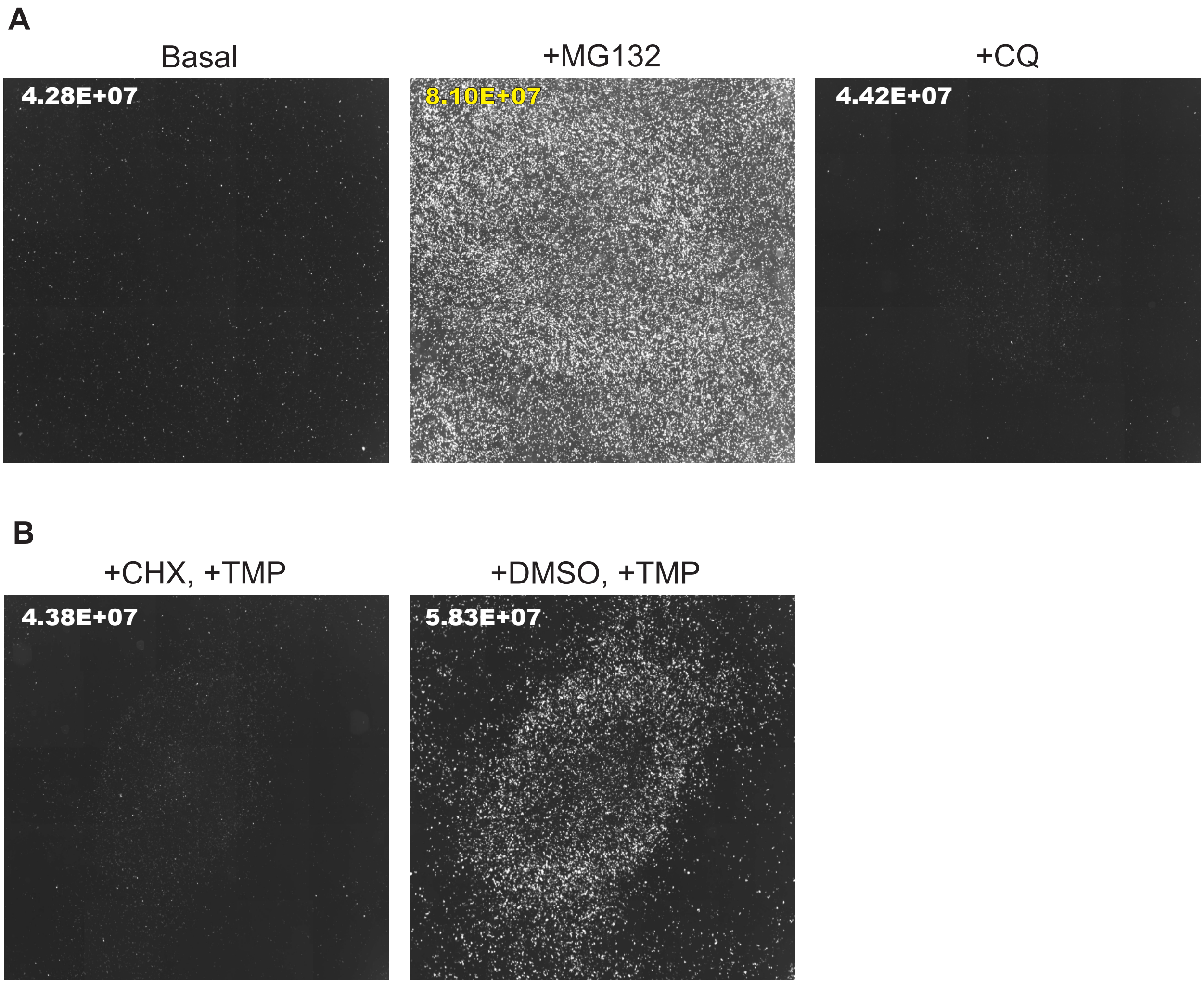

Fig. S3

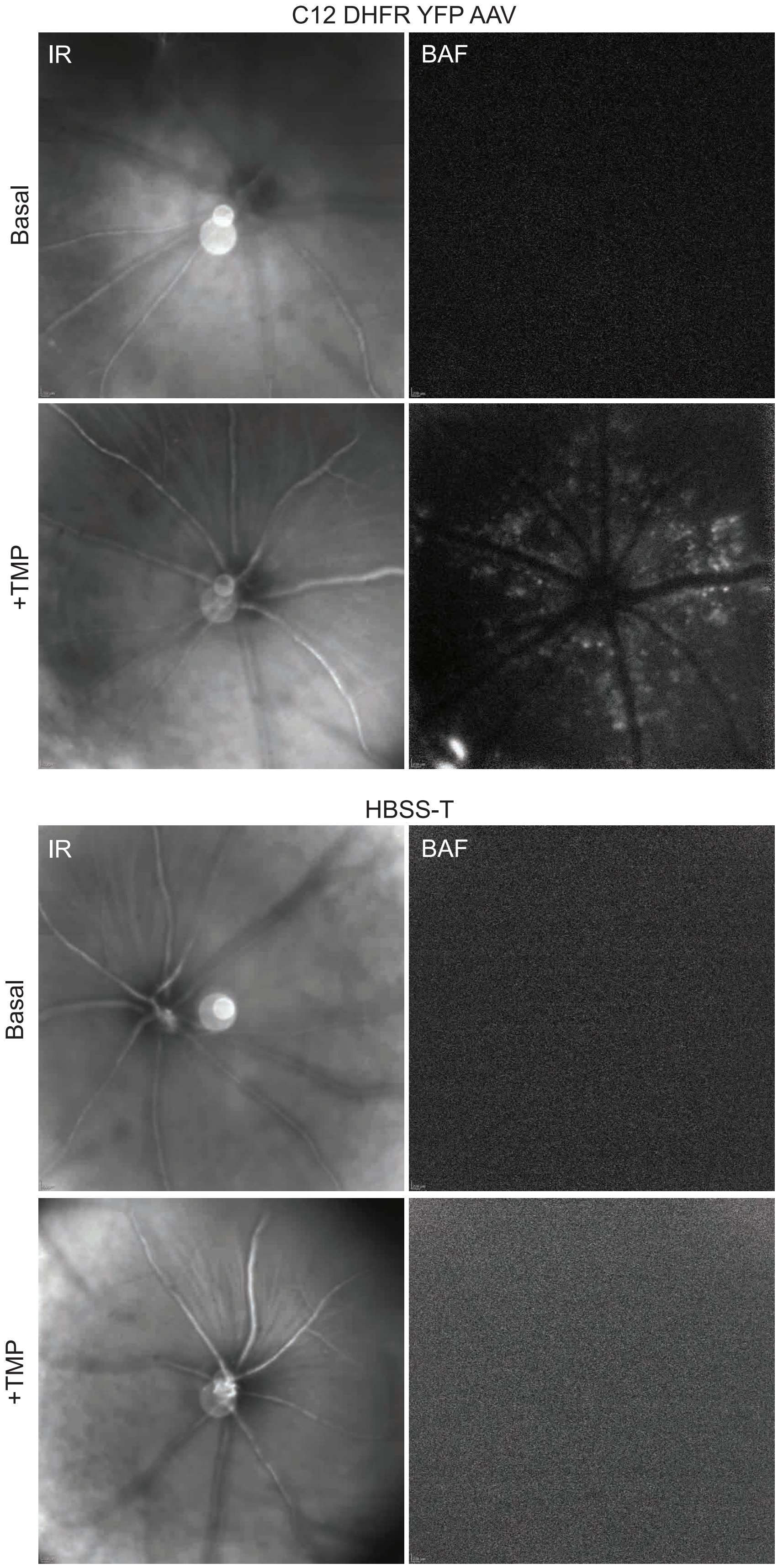

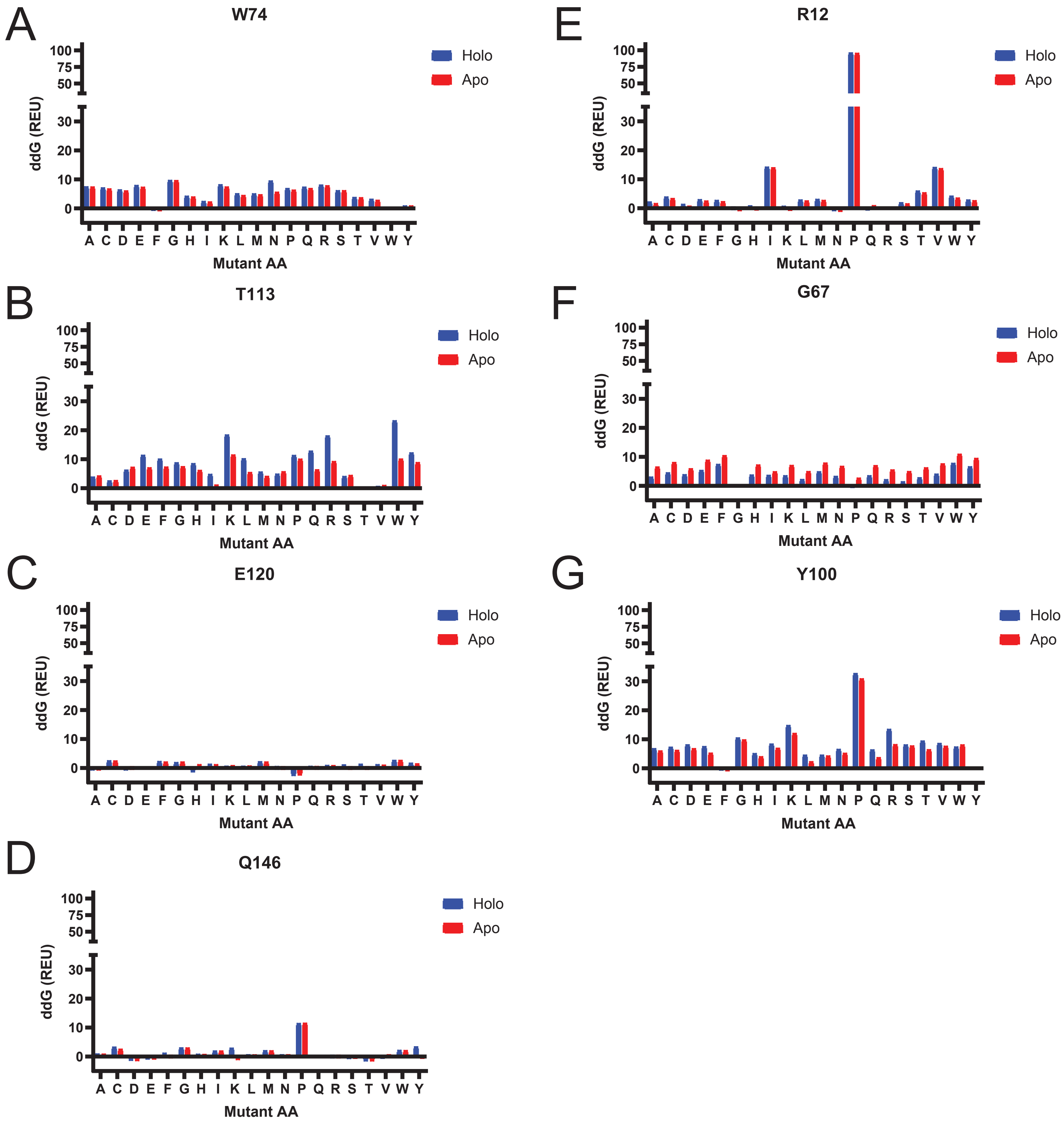

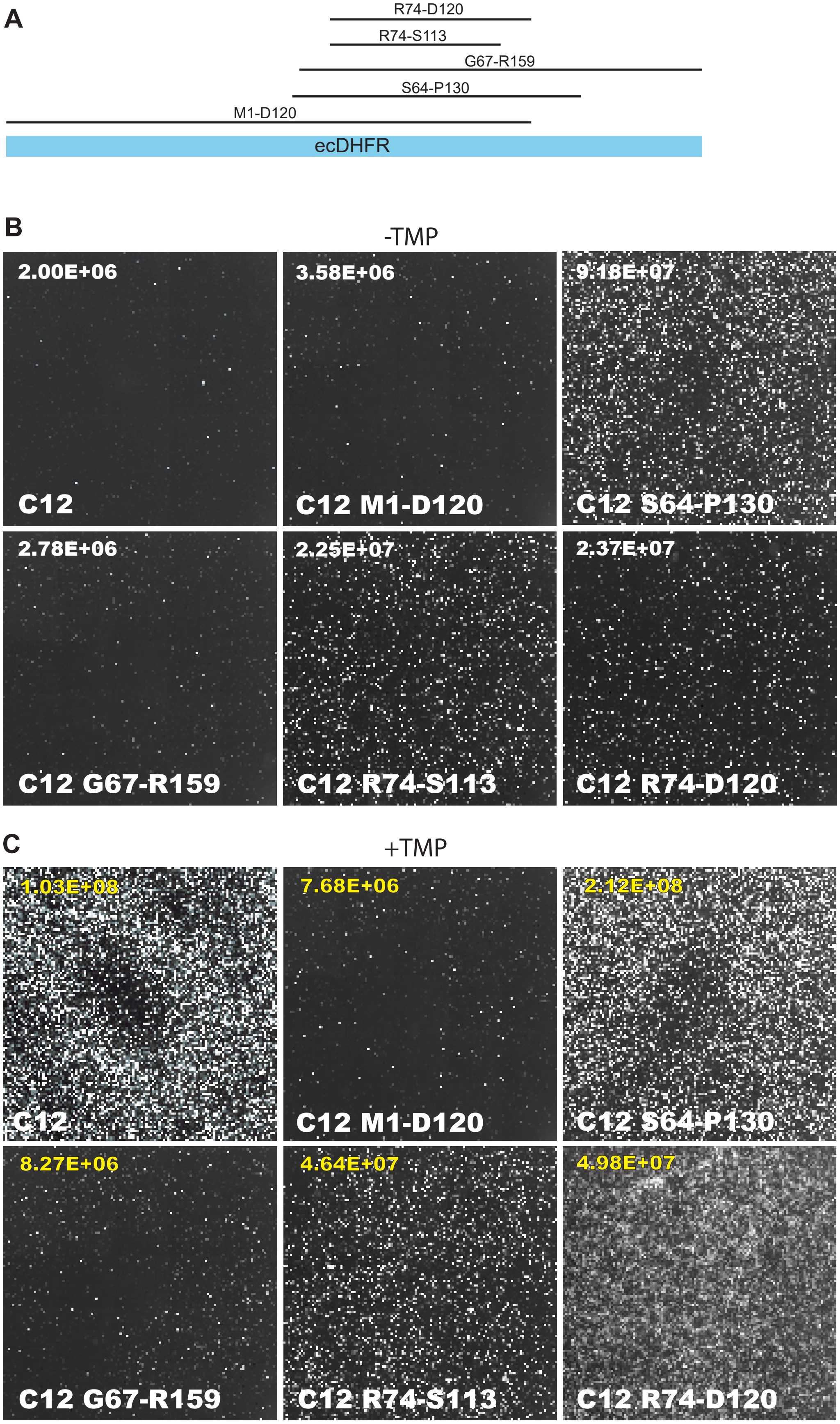
